# BSG/CD147 and ACE2 receptors facilitate SARS-CoV-2 infection of human iPS cell-derived kidney podocytes

**DOI:** 10.1101/2021.11.16.468893

**Authors:** Titilola D. Kalejaiye, Rohan Bhattacharya, Morgan A. Burt, Tatianna Travieso, Arinze E. Okafor, Xingrui Mou, Maria Blasi, Samira Musah

## Abstract

**Background:** Severe acute respiratory syndrome coronavirus 2 (SARS-CoV-2) causes the Coronavirus disease 2019 (COVID-19), which was declared a pandemic by the World Health Organization (WHO) in March 2020. The disease has caused more than 5.1 million deaths worldwide. While cells in the respiratory system are frequently the initial target for SARS-CoV-2, clinical studies suggest that COVID-19 can become a multi-organ disease in the most severe cases. Still, the direct affinity of SARS-CoV-2 for cells in other organs such as the kidneys, which are often affected in severe COVID-19, remains poorly understood.

**Method:** In this study, we employed a human induced pluripotent stem (iPS) cell-derived model to investigate the affinity of SARS-CoV-2 for kidney glomerular podocytes. We studied uptake of the live SARS-CoV-2 virus as well as pseudotyped viral particles by human iPS cell derived podocytes using qPCR, western blot, and immunofluorescence. Global gene expression and qPCR analyses revealed that human iPS cell-derived podocytes express many host factor genes (including ACE2, BSG/CD147, PLS3, ACTR3, DOCK7, TMPRSS2, CTSL CD209, and CD33) associated with SARS-CoV-2 binding and viral processing.

**Result:** Infection of podocytes with live SARS-CoV-2 or spike-pseudotyped lentiviral particles revealed viral uptake by the cells at low Multiplicity of Infection (MOI of 0.01) as confirmed by RNA quantification and immunofluorescence studies. Our results also indicate that direct infection of human iPS cell-derived podocytes by SARS-CoV-2 virus can cause cell death and podocyte foot process retraction, a hallmark of podocytopathies and progressive glomerular diseases including collapsing glomerulopathy observed in patients with severe COVID-19 disease. Additionally, antibody blocking experiments identified BSG/CD147 and ACE2 receptors as key mediators of spike binding activity in human iPS cell-derived podocytes.

**Conclusion:** These results show that SARS-CoV-2 can infect kidney glomerular podocytes *in vitro*. These results also show that the uptake of SARS-CoV-2 by kidney podocytes occurs via multiple binding interactions and partners, which may underlie the high affinity of SARS-CoV-2 for kidney tissues. This stem cell-derived model is potentially useful for kidney-specific antiviral drug screening and mechanistic studies of COVID-19 organotropism.

**Significant statement:** Many patients with COVID19 disease exhibit multiorgan complications, suggesting that SARS-CoV-2 infection can extend beyond the respiratory system. Acute kidney injury is a common COVID-19 complication contributing to increased morbidity and mortality. Still, SARS-Cov-2 affinity for specialized kidney cells remain less clear. By leveraging our protocol for stem cell differentiation, we show that SARS-CoV-2 can directly infect kidney glomerular podocytes by using multiple Spike-binding proteins including ACE2 and BSG/CD147. Our results also indicate that infection by SARS-CoV-2 virus can cause podocyte cell death and foot process effacement, a hallmark of podocytopathies including collapsing glomerulopathy observed in patients with severe COVID-19 disease. This stem cell-derived model is potentially useful for kidney-specific antiviral drug screening and mechanistic studies of COVID-19 organotropism.

## Introduction

In March 2020, Coronavirus disease 2019 (COVID-19) - caused by the severe acute respiratory syndrome coronavirus 2 (SARS-CoV-2) - was declared a pandemic by the World Health Organization (WHO).^1, 2^ Since then, the disease has affected more than 253 million people globally and caused over 5.1 million deaths worldwide^3^). Coronaviruses are positive-strand RNA viruses (+RNA)^4^ with large genomes^5–7^ that are packaged into a helical nucleocapsid and encode for structural protein genes including Spike protein (S), Envelope protein (E), Membrane (M) and Nucleocapsid (N) ^8^ along with several accessory proteins.^9^ The spike protein facilitates entry of the virus (i.e. receptor binding and membrane fusion) into target cells.^10–12^ Notably, the S-glycoprotein is the target of neutralizing antibodies and for the development of vaccines^13^ since it is the primary antigenic determinant of infectivity and pathogenicity.^14^

The extent of SARS-CoV-2 tropism (cell types or tissues permissive to viral infection) is determined by the expression of a suitable receptor on the cell surface as well as the presence of a host-encoded protease proximally positioned to the site of receptor binding to enable cleavage of the S protein for viral processing.^9, 15^ Angiotensin-Converting Enzyme 2 (ACE2) is recognized as the main receptor for SARS-CoV-2 binding to cells and tissues.^1, 6, 10, 12, 16^ On the other hand, Basigin (BSG, also known as CD147 or EMMPRIN), a transmembrane glycoprotein has been shown to be an alternative route for SARS-CoV and SARS-CoV-2 invasion of host cells.^17, 18^ For instance, BSG/CD147 is shown to interact with S protein *in vitro* and facilitate entry of SARS-CoV and SARS-CoV-2 in Vero and HEK 293T cell.^18, 19^ Additionally, BSG/CD147 was shown to allow entry of SARS-CoV-2 into baby hamster kidney cells (BHK-21) but only when they overexpress human CD147 (BHK-21-CD147 cells). Similarly viral load was detected in the lungs of SARS-CoV-2 infected mice expressing human CD147 (hCD147).^18^ Once spike binds to its receptor,^13^ the activity of proteases such as Transmembrane Serine Protease 2 (TMPRSS2) or cathepsin L (CTSL) promote fusion and internalization of the receptor-viral spike complex.^20^

Although SARS-CoV-2 primarily infects cells in the respiratory tract, other tissues and organs can also be vulnerable to the virus and result in a broad array of complications in the renal, cardiovascular, gastrointestinal and nervous systems.^21–23^ Several *in vitro* studies have examined the impact of SARS-CoV-2 infection in lung and cardiac cells.^24–27^ SARS-CoV-2 viral RNA has been detected in multiple organs including the kidneys.^28, 29^ Intriguingly, acute kidney injury and cardiac injury are common in COVID-19 patients^15, 30, 31^ and have been associated with increased morbidity and mortality.^32, 33^ However, it remains unclear whether the virus can directly infect specific types of human kidney cells such as podocytes, a specialized epithelial cell type often targeted in many forms of kidney diseases. Additionally, collapsing glomerulopathy has been reported in COVID-19 patients and the phenomenom has been termed COVID-19-associated nephropathy (COVAN).^34^ COVAN resembles human immunodeficiency virus (HIV) –associated nephropathy (HIVAN), a kidney disease caused by HIV infection^35^ and resulting in CKD and kidney failure.^36^ Notably, collapsing glomerulopathy constitutes a new renal manifestation of COVID-19 that may also arise from genetic predisposition.^37^

One of the most common clinical indications of COVID-19 is reduced blood lymphocyte count which can significantly increase virulence.^1^ This eventually leads to cytokine release syndrome (CRS) which is highly associated with respiratory failure, acute respiratory distress syndrome (ARDS) and other adverse clinical consequences.^38^ Once the lungs become infected, the virus moves through the blood^39^ to other organs such as the kidneys where it can cause damage to resident renal cells.^30^ The exact mechanism of kidney pathology in COVID-19 is not clear, although there are suggestions that it is either through virus-activated cytokine storm due to sepsis or direct injury on kidney cells caused by the virus.^22^

Within the kidney, podocytes and proximal tubules play important roles in renal filtration, reabsorption and excretion.^40^ The glomerulus, a network of capillaries, is the primary site for blood filtration. The glomerular filtration barrier consist of interdigitated podocytes separated from fenestrated glomerular endothelial cells by the glomerular basement membrane (GBM), all of which function together to facilitate the selective filtration of toxins and waste from the blood.^41–43^ Importantly, podocytes are particularly vulnerable to bacterial and viral attacks and injury which can result in retraction of podocyte foot processes and effacement, causing abnormal leakage of proteins into the urine (proteinuria).^44^ Given that SARS-CoV-2 has been found in nephrin-positive cells of the kidneys of COVID-19 patients,^29^ we hypothesized that podocytes could be direct targets for SARS-CoV-2 infection.

Understanding the susceptibility of organ-specific cell types to SARS-CoV-2 infection and COVID-19 disease mechanisms rely on the availability of robust experimental models that can closely mimic the functional phenotype and developmental status of human cells and tissues. Human pluripotent stem cells provide an enormous advantage for such studies due to their ability to proliferate indefinitely and differentiate into organ-specific cell types. Thus, derivatives of human pluripotent stem cells can be used for disease modelling and drug discovery assays.^45, 46^ Stem cells can be differentiated into kidney organoids ^47–50^ or even more specialized cells such as podocytes.^43^ Monteil et al. studied SARS-CoV-2 infectivity in induced pluripotent stem cells derived kidney organoids using a protocol developed by Montserrat and colleagues.^47, 49^ However, organoids mostly generate immature cells that resemble fetal stages of development^51^ and lack the functional maturation and phenotype^52, 53^ necessary for studying postnatal kidney biology and disease mechanisms. We previously developed a method for the direct differentiation of human iPS cells into cells that exhibit morphological, molecular, and functional characteristics of the mature human kidney glomerular podocytes.^43, 54, 55^ We employed this model to study the susceptibility of kidney podocytes to SARS-CoV-2 infection. Furthermore, we quantified the basal mRNA expression of key SARS-CoV-2 and coronavirus-associated receptors and factors^15^ and found that podocytes express many of the genes involved in viral entry and processing. We confirmed protein-level expression of some of these factors using western blot and immunocytochemistry analyses. We also uncovered a role for BSG/CD147 as a receptor for SARS-CoV-2 viral entry into podocytes. Together, our results demonstrate for the first time that specialized human iPS cell-derived podocytes can be directly infected by SARS-CoV-2 at low multiplicity of infection (MOI) through ACE2 and BSG/CD147 receptors, resulting in loss of arborized foot-like processes as well as changes in the expression of genes important for cell lineage determination and function.

## Materials and Methods

### Cell culture conditions

All cell lines used for this study were obtained under appropriate material transfer agreements and approved by all involved institutional review boards. All cells were tested for and shown to be devoid of mycoplasma contamination (Mycoplasma PCR Detection Kit from abm, G238). Human colon epithelial (Caco-2) (ATCC, HTB-37) and human embryonic kidney (HEK 293T) (ATCC, CRL-3216) cell lines were cultured in high-glucose Dulbecco’s Modified Eagle Medium (DMEM; Gibco, 12634010) media supplemented with 10% fetal bovine serum (FBS; Gibco; 10082147) with L-Glutamine (Gibco; 25030081) and 1X Penicillin/Streptomycin (Gibco; 15140122). HEK 293T cells were split (1:10) while Caco-2 cells were split (1:5) every 3 days. Human lung (Calu-3) (ATCC, HTB-55) cells were cultured in Minimum Essential Media (MEM) (Gibco; 11095080) supplemented with 10% FBS with 1mM sodium pyruvate (Gibco; 11360070), MEM non-essential amino acids (NEAA) (Gibco; 11140050) with 1X Penicillin/Streptomycin. Human induced pluripotent stem (Human iPS) cell line used for this study (PGP1 – the Personal Genome Project ^56^) were tested and shown to be free of mycoplasma contamination. The cell line had normal karyotype. Human iPS cells were cultured in mTeSR1 (StemCell Technologies; 85870) medium without antibiotics and split (1:6) every 4 to 5 days. All cells were incubated in a 37 °C incubator with 5% CO_2_.

### Differentiation of human iPS cells into podocytes

Mature human glomerular podocytes were generated using previously published protocol ^43, 54, 55^. Briefly, human induced pluripotent stem (iPS) cells cultured on Matrigel-coated plates were dissociated with warm enzyme-free dissociation buffer (Gibco, 13150-016) and centrifuged twice at 200x*g* for 5 min each in advanced DMEM/F12 (Gibco; 12634010). The DMEM/F12 was aspirated off and the cells were resuspended in mesoderm induction media (consisting of DMEM/F12 with GlutaMax (Gibco; 10565042) supplemented with 100 ng/ml activin A (Invitrogen; PHC9564), 3 μM CHIR99021 (Stemgent; 04-0004), 10 μM Y27632 (TOCRIS; 1254) and 1X B27 serum-free supplement (Gibco; 17504044) and plated at a seeding density of 100,000 cells per well of a 12-well plate. The cells were cultured in the mesoderm induction medium for 2 days with daily medium change and after two days, intermediate mesoderm differentiation was initiated by feeding the cells with intermediate mesoderm induction medium (containing DMEM/F12 with GlutaMax supplemented with 100 ng/ml BMP7 (Invitrogen; Phc9543), 3 μM CHIR99021 and 1X B27 serum-free supplement) for a minimum of 14 days. Podocyte induction was initiated by dissociating the intermediate mesoderm cells with 0.05% trysin-EDTA (Gibco; 25300-054) for 5 min with subsequent quenching of the enzyme with 10% FBS in DMEM/F12 (trypsin neutralizing solution). Adhered cells were gently scrapped using a cell scrapper and then pipetted using a P1000 barrier tip to obtain individualized cells. The cell suspension was transferred into a 50 ml falcon tube containing 30 ml DMEM/F12 and centrifuged twice at 200x*g* for 5 min each. The cell pellet was resuspended in podocyte induction mediaand plated on a freshly prepared laminin-511-E8-coated plates at a seeding density of 100,000 cells per well of a 12 well plate. The cells were fed podocyte induction media containing advanced DMEM/F12 with GlutaMax supplemented with 100 ng/ml BMP7, 100 ng/ml activin A, 50 ng/ml VEGF (Gibco; PHC9394), 3 μM CHIR99021, 1X B27 serum-free supplement, and 0.1 μM all-trans retinoic acid (Stem Cell Technologies; 72262) for 5 days. Mature podocytes were maintained in CultureBoost-R (Cell Systems; 4Z0-500).

### Production of SARS-CoV-2 S-pseudotyped lentiviral particles

The S-pseudotyped lentiviral particles were generated by transfecting HEK 293T cells as illustrated in **Figure 1C**. Briefly, HEK 293T cells were seeded in DMEM-10 growth media in 75 cm^3^ flask until about 65 to 75% confluent. The cells were then transfected with the plasmids required for the lentiviral production using Lipofectamine 3000 reagent (Invitrogen, L3000015) in Opti-MEM (Gibco; 31985070) following manufacturer’s instructions. Briefly, 20 µg of plasmid DNA (total plasmid mix) per flask was mixed in Opti-MEM and P3000 reagent and left at room temperature for 5 minutes. Plasmids mix for each transfection consisted of psPax2 (packaging; Addgene plasmid # 12260), pCMV-SCoV-2S (Spike envelope plasmid; Sinobiologicals – Cat #: VG40589-UT) and pLJM1-EGFP (reporter; Addgene plasmid #19319) in a ratio of 1:1:2 respectively. After 5 min incubation, the plasmid DNA mix in Opti-MEM-P3000 media was then mixed with the transfection reagent (Opti-MEM and Lipofectamine 3000 reagent) and incubated at room temperature for 10-15 minutes. Appropriate volumes of transfection mixture was used to transfect HEK 293Tcells in each flask and incubated in a 37 °C incubator with 5% CO2 for 6 hours. Lentivirus pseudotyped with the vesicular stomatitis virus spike G (pCMV-VSV-G; Addgene plasmid # 8454) was used as a positive control and the bald lentivirus having no coat was used as a negative control.

**Figure 1:**
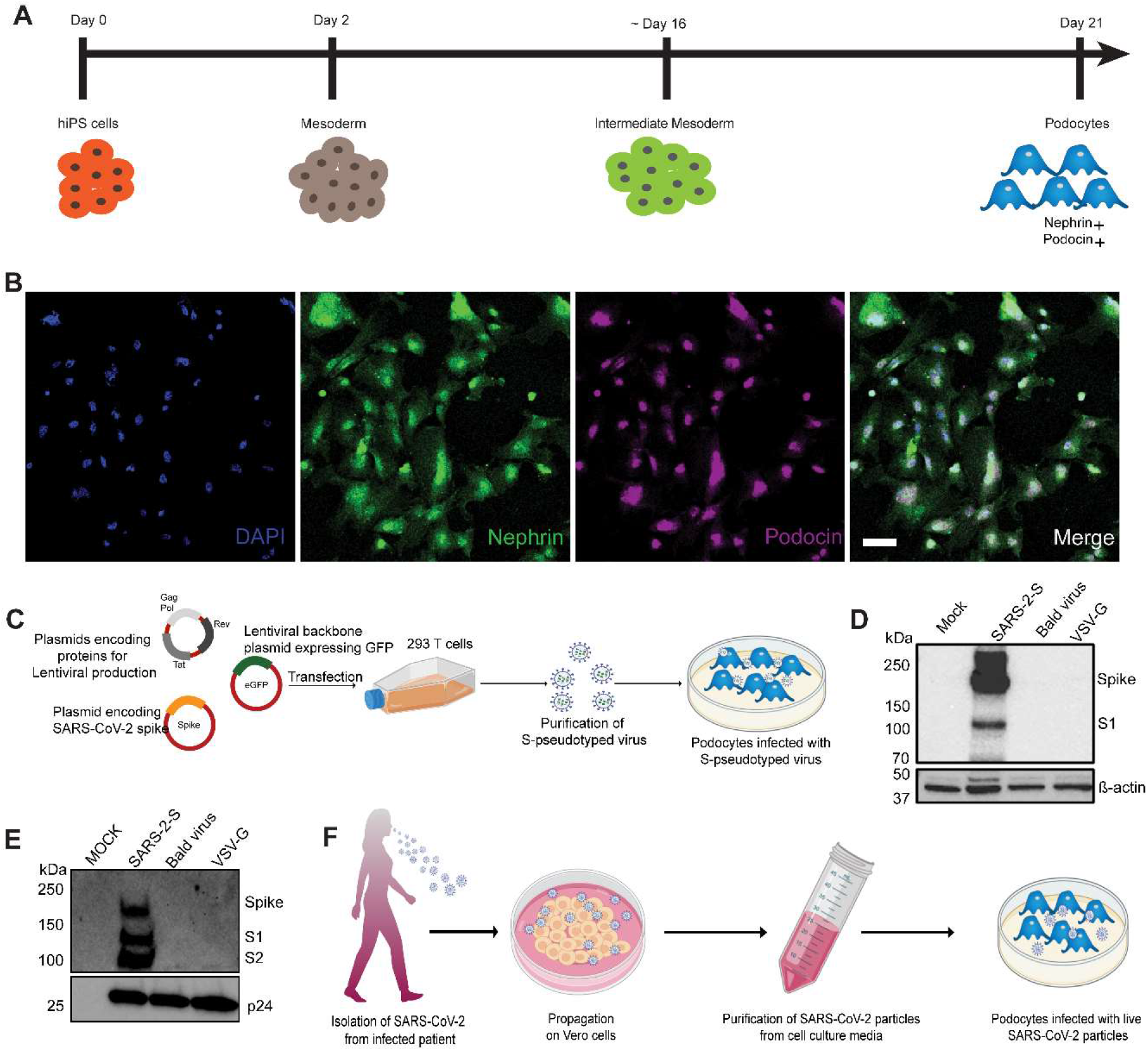
Establishment of a method to examine susceptibility of human iPS cell-derived podocytes to SARS-CoV-2 infection. (**A**) Schematic overview of the protocol for the differentiation of mature podocytes from human iPS cells; adapted from ^43^. (**B**) Human iPS cell-derived podocytes express the lineage specific markers nephrin (green) and podocin (magenta). Cells were counterstained with DAPI (blue) nuclear marker. Scale bar, 100 μm. (**C**) Schematic depicting the lentiviral vectors used to produce S-pseudotyped virus from 293T cells and infection of human iPS cell-derived podocytes *in vitro*. (**D-E**) Western blot confirming the successful transfection of 293T cells and production of S-pseudotyped virus indicated by the presence of Spike protein (190kDA) and its cleavage products S1 (∼110kDa) in both (**D**) the cell lysates (β-actin used as loading control) and (**E**) purified viral particle (normalized to HIV viral protein p24) with extra band for S2 (∼100kDa). Mock for figure D represents lysate from untransfected cells while in E, mock represents pseudoviral particle dissolving media (TBS); SARS-2-S represent lysate from cells transfected with spike expressing plasmid or S-pseudotyped particle; Bald virus represent lysate from cells transfected with no envelope plasmid or the resulting pseudovirus particle without viral envelope protein; VSV-G represent lysate from cells transfected with plasmid expressing VSV-G or pseudotyped particle with VSV-G envelope (**F**) Schematic showing propagation of patient-derived SARS-CoV-2 virus in Vero E6 cells followed by purification and infection of human iPS cell-derived podocytes *in vitro*.

At 6 hours post-transfection, the culture medium was replaced with fresh pre-warmed DMEM-10. After an additional 24 hours (30 hours post-transfection) and 48 hours (54 hours post-transfection), the lentiviral particles were harvested by collecting the supernatant from each flask, centrifuging at 1000x*g* for 5 minutes and filtering through a 0.45 µm SFCA low protein-binding filter. Samples were then subjected to ultracentrifugation over a 28% sucrose cushion (Sucrose/PBS; Sigma S7903-1KG) at 100,000x*g* for 3 hours at 4 °C. The pellet was resuspended in 1X Tris buffered saline (TBS, Bio-Rad; 1706345), and then aliquoted and stored at −80 °C to avoid repeated freeze-thaw cycles.

### Infection of cells with S-pseudotyped virus

Cells were cultured in the appropriate culture media and infected with S-pseudotyped virus in the presence of polybrene (Sigma; TR-1003) to a final concentration of 5 µg/ml.

### SARS-CoV-2 expansion in Vero E6 cells and titration

All experiments with the live virus were performed under Biosafety Level 3 (BSL-3) in the Duke Regional Biocontainment Laboratory at the Duke Human Vaccine Institute (DHVI) in compliance with the BSL-3 laboratory safety protocols and guidelines from the CDC for handling SARS-CoV-2.

SARS-CoV-2 USA-WA1/2020 (BEI Resources; NR-52281) was propagated in Vero E6 cells at a MOI of 0.001 in DMEM supplemented with 2% FBS, 1X Penicillin/Streptomycin, 1 mM Sodium pyruvate and 1X Non-Essential Amino Acid (NEAA) at 37 °C in 5% CO_2_. Four days post infection (pi), supernatant containing the released virus were harvested, centrifuged at 1500 rpm for 5 minutes and filtered through a 0.22 µM filter. Samples were aliquoted and stored at −80 °C until further use.

Plaque assay was done to determine the titer of the viral stock. Briefly, 0.72 × 10^6^ Vero E6 cells were seeded in 24 well plates. The virus stock was diluted serially (10-fold), and the dilutions were used to infect monolayer of Vero E6 cells at 37 °C in 5% CO_2_. After an hour of incubation, cells were overlayed with media containing carboxy-methyl cellulose (CMC) (0.6% CMC), MEM supplemented with 2% fetal bovine serum (FBS), 1mM sodium pyruvate (Gibco), 1X NEAA (Gibco), 0.3% sodium bicarbonate (Gibco) and 1X GlutaMAX (Gibco) with 1X Penicillin/Streptomycin. After 4 days of incubation at 37 °C in 5% CO_2_, cells were stained stained with 1% crystal violet in 10% neutral buffered formalin (NBF) and the number of plaque forming units per ml (pfu/ml) was determined.

### Infection of human iPS cell-derived podocytes with live virus

1 × 10^5^ intermediate mesoderm cells were differentiated to podocytes (per well of a 12 well plate). After 5 days of induction, podocytes were incubated with the SARS-CoV-2 virus at an MOI 0.01, 0.1 or 1 at 37 °C and 5% CO_2_ with intermittent plate rocking. To obtain the desired MOI, SARS-CoV-2 was diluted in CultureBoost-R and incubated with the podocytes for 1 hour at 37 °C. After 1 hour of incubation, the virus-containing supernatant was aspirated, and cells were washed twice with 1X PBS. Fresh maintenance medium was then added, and cells incubated for either 24, 48 or 72 hours at 37 °C and 5% CO_2_. Uninfected controls were incubated with CultureBoost-R only.

### Infectious viral titer of supernatant

Cellular supernatant was collected from podocytes infected with SARS-CoV-2 virus at MOI of 0.01, 0.1 and 1, at 24hrs, 48hrs and 72hrs post infection. The supernatant was clarified by centrifugation at 1500 rpm for 5 minutes and the infectious viral titer was measured by plaque assay as described above.

### qRT-PCR for detection of intracellular and cell-free viral RNA

SARS-CoV-2 RNA was extracted from the supernatant or cell pellet of infected podocytes using the QIAamp viral RNA mini kit (Qiagen; cat # 52904). qRT-PCR was performed with primers specific for target genes (see **Supplementary Table 1** for the list of primers) using the Luna universal One-Step RT-qPCR kit (NEB; E3005). Experiment was performed using the QuantStudio3 (Applied Biosystems) with the following thermal cycling steps; 55 °C for 10 minutes, 95 °C for 1 minute and 40 cycles of 95 °C for 10 seconds and 60 °C for 1 minute according to manufacturer’s protocol.

### qRT-PCR and qPCR analysis of infected cells

Cell pellets were washed and lysed using RA1 RNA extraction buffer and purified using the NucleoSpin RNA kit (MAcherey-Nagel; 740955.250) following the manufacturer’s instructions. RNA from infected and control podocytes were harvested using NucleoSpin RNA kit. The RNA was quantified by nanodrop (Thermo Fisher). 0.5 µg to 1 µg of RNA was converted to cDNA for qPCR. cDNA synthesis was done using SuperScript III Reverse Transcriptase (Invitrogen; 18080-085) and qPCR was performed using qPCR SYBR Master Mix (Promega; A6001). Quantitative PCR was performed with QuantStudio3 (Applied Biosystems) using the thermal cycling steps; 50 °C for 2 minutes, 95 °C for 10 minutes and 40 cycles of 95 °C for 15 seconds and 60 °C for 1 minute. Delta cycle threshold (ΔCt) was determined relative to GAPDH. Viral RNA from pseudovirus infected cells was also quantified by qRT-PCR using the Lenti-X qRT-PCR titration kit (Clontech; 631235) following manufacturer’s instruction. Primer sequences are provided in the **Supplementary Table 1**.

### Western blot analyses

For western blotting, cells were first lysed using RIPA buffer (Sigma; R0278-500ML) supplemented with protease inhibitor cocktail (Roche) at 4 °C with shaking for 30 min for protein extraction. Pierce BCA protein assay Kit (ThermoF Scientific; 23227) was used for protein quantification. 15 µg of the extracted protein samples were boiled for 5 minutes at 95 °C in 4X laemlli buffer (BioRad; 1610747), run on mini-PROTEAN TGX precast gels (Bio-Rad; 4568083) and then transferred to PVDF membrane blot (Bio-Rad; 1620175). The blots were blocked in 5% non-fat milk made in TBS-T (50 mM Tris-HCl, 150 mM NaCl, 0.1% Tween-20) for 1 hour and incubated with the primary antibodies (dilutions in the key resource table) in blocking buffer overnight at 4 °C. The next day, horseradish-peroxidase-conjugated rabbit anti-goat (R&D Systems; HAF017), goat anti-rabbit (CST; 7074) or goat anti-mouse (CST; 7076) antibody was added, and the blot was incubated for 1 hour at room temperature. The membranes were developed with the SuperSignal West Femto substrate (Thermo Fisher Scientific, 34095) by following manufacturer’s protocol. The chemi-luminiscent signals were acquired using a GelDoc Imager (Bio-Rad).

### Identification of spike associated host factors expressed by podocyte

To identify podocyte host factors that could facilitate viral entry and replication, we integrated the BioGRID interaction database with transcriptomic data previously generated in our lab. BioGRID is an expansive database of experimentally verified protein-protein and genetic interactions as assembled and curated from tens of thousands of studies.^57^ Firstly the latest release of the BioGRID interaction database (as at when this study was carried out) for coronaviruses was downloaded from the archive at https://downloads.thebiogrid.org/Download/BioGRID/Release-Archive/BIOGRID-3.5.188/BIOGRID-CORONAVIRUS-3.5.188.tab3.zip. The interaction network file was then opened using Cytoscape (v3.8.0) and filtered to obtain only edges linking human proteins to the SARS-CoV2 spike protein.

We extracted podocyte gene expression data for spike-binding proteins and integrated it with the network table obtained from BioGRID. The microarray transcriptomic data for human iPS cell-derived podocytes used in this 9tudy had been generated in a previous study.^54^ The podocyte microarray gene expression data were analyzed using standard pipeline. Briefly, the raw expression data were normalized by robust multiarray averaging ^58^ and the Human Gene 2.0 ST Affymetrix array mapping obtained from the ENSEMBL mart database was used to map probe IDs to gene IDs. The podocyte transcriptomic data was analyzed using the Bioconductor packages, oligo (v3.11), biomaRt, and pd.hugene.2.0.st.^59^ The expression data for these proteins were then used to annotate a network visualization of these interactions on Cytoscape.

### Immunofluorescence Imaging

For immunofluorescent imaging, human iPS cell-derived podocytes (infected and control) were fixed with 4% paraformaldehyde (PFA) in PBS for 20 to 30 minutes at room temperature and permeabilized using 0.125% Triton X-100 (Sigma-Aldrich) in PBS for 5 minutes. Cells were blocked with 1% BSA/PBS-T for 30 minutes at room temperature and then incubated with primary antibody diluted in the blocking buffer overnight at 4 °C. After overnight incubation, cells were incubated with Alexa Fluor-488 or Alexa Fluor-594 donkey (Invitrogen, 1:1000) secondary antibodies diluted in blocking buffer for 1 hour at room temperature. Cells were afterwards counterstained with 4′,6-diamidino-2-phenylindole (DAPI, Invitrogen). The primary antibodies used were nephrin (Progen, GP-N2); podocin (Abcam, ab50339); anti-SARS-CoV2 spike (ProSci, 3525); anti-SARS-CoV-2 N protein (Sinobiological, 40143-R019); VSV-G (Santa Cruz Biotechnology, sc-365019); Human/Mouse/Rat/Hamster ACE-2 (R&D systems, AF933), Human TRA-1-85/CD147 (R&D systems, MAB3195), Cathepsin L (Santa Cruz Biotechnology, sc-32320), TMPRSS2 (Santa Cruz Biotechnology, sc-515727) and DC-SIGN/CD209 (Santa Cruz Biotechnology, sc-65740). Images were acquired using an M7000 epifluorescence microscope (Invitrogen, AMF7000) equipped with 10x/0.30 LWDPH with 7.13 mm WD and 20x/0.45 LWDPH with 6.12 mm WD objectives. Confocal images were captured using a Zeiss 880 inverted confocal Airyscan with a 10x/0.30 EC Plan-Neofluar air lens with 5.2 mm objectiveat the Duke Light Microscopy Core Facility.

### Blocking of ACE2 and BSG/CD147 protein with antibodies

For the blocking of ACE2 and/or BSG/CD147 epitope, we infected podocytes with the S-pseudotyped lentivirus. Approximately 1.5 hours prior to infecting cells, antibody dilutions were prepared in the CultureBoost-R. We performed the blocking experiment using an ACE2 polyclonal goat antibody (Cat # AF933; R&D systems) and CD147 (BSG) mouse monoclonal antibody (Human TRA-1-85/CD147 MAb (Clone TRA-1-85)-MAB3159; R&D systems). The human iPS cell-induced podocytes were pre-treated with serial dilutions of ACE2 antibody, BSG/CD147 antibody or both for 1 hour. Unblocked cells and uninfected (mock) cells were used as control. After 1 hour of incubation, the pseudoviral particles (MOI-0.02) were added to each well and incubated for 48 to 60 hours. After 60 hours, cells were washed and lysed in RNA extraction buffer. RNA was purified using the Macherey Nagel RNA extraction kit following manufacturer’s instruction and viral RNA uptake was quantified using the Luna universal One-Step RT-qPCR kit (NEB; E3005).

### Quantification and Statistical analysis

All experiments were done in 3 independent biological replicates unless otherwise indicated. N=3. One-way analysis of variance (ANOVA) with Šidák’s posttest or multiple t-test was used to test for statistical significance. Only p values of 0.05 or lower were considered statistically significant (p > 0.05 [ns, not significant], p < 0.05 [*], p < 0.01 [**], p < 0.001 [***], p < 0.0001 [****]). For all statistical analyses, the GraphPad Prism 9 software package was used (GraphPad Software).

## Results

### Generation of mature kidney glomerular podocytes from human iPS cells

We performed directed differentiation of human iPS cells to generate glomerular podocytes by following our previously described protocol.^43, 54, 55^ The differentiation of human iPS cells to glomerular podocytes was performed in three stages using defined media (**Figure 1A**). Briefly, human iPS cells were first differentiated into mesoderm by incubation for two days in a mesoderm induction medium. The resulting mesoderm cells were then incubated with an intermediate mesoderm (IM) induction medium for 14 days. Subsequent culture of the IM cells in podocyte induction medium for 5 days enabled the derivation of mature podocytes that exhibited highly specialized morphological features and expressed podocyte-specific markers including nephrin and podocin (**Figure 1B**). Notably, our methodology enables high-yield derivation of podocytes with mature phenotype and functional characteristics including the formation of an intact glomerular filtration barrier and selective removal of molecules from the vasculature^43^ – thus, avoiding the limitations of fetal-like cells often produced by organoid models.^20, 49^

In this work, we used the live virus (SARS-CoV-2 USA-WA1/2020 - BEI Resources; NR-52281) and an in-house produced S-pseudotyped virus (details below) to study the effect of SARS-CoV-2 on human iPS cell-derived podocytes.

### Podocytes are permissive to S-pseudotyped viral infection

The spike surface envelope glycoprotein (S) facilitates binding and entry of coronavirus including SARS-CoV and SARS-CoV-2 into cells,^13^ and it exhibits capabilities for receptor binding and membrane fusion.^60, 61^ We initially employed S-pseudotyped virus to study viral entry and uptake into podocytes. To generate the S-pseudotyped virus, we used an HIV-1-based S-pseudotyped lentiviral vector as illustrated in **Figure 1C**. HEK 293T (293T) cells were co-transfected with lentiviral vector plasmids required to generate the S-pseudotyped virus, which included a plasmid encoding a green fluorescent reporter protein, a plasmid expressing HIV proteins (gag, tat and pol) necessary for viral assembly and a SARS-CoV-2-S expressing plasmid (**Figure 1C**). To generate control pseudotyped viruses, we used the vesicular stromatitis virus glycoprotein (VSV-G; control envelope) plasmid and a ‘bald’ virus lacking an envelope protein. Western blot analysis of 293T cell lysates confirmed the presence of spike protein in only the cells that were transfected with the spike plasmid and not in the cell lysates obtained from VSV-G or bald virus transfection **(Figure 1D**). Western blot analysis of the 293T cell supernatant from S-pseudotyped particle produced three major bands at 190, 110, and 100 kDa showing the presence of full-length and cleaved S proteins (S1 and S2) respectively, as well as the HIV gag protein (p24). These results confirmed the incorporation of the S protein in the pseudoviral particles and the successful generation of S-pseudotyped virus with the SARS-CoV-2 spike protein (SARS-2-S) (**Figure 1E**). As expected, the VSV-G and bald pseudoviruses produced only a band for HIV p24 while no band was detected in the mock (medium control) (**Figure 1E**). Live virus Infection of human iPS cell-derived podocytes was performed using SARS-CoV-2 strain USA-WA1/2020 grown on Vero E6 cells as illustrated in **Figure 1F**.

We initially inoculated human iPS cell-derived podocytes with the S-pseudotyped virus to examine their permissiveness to the virus. As changes in inoculum volume and transduction time can influence intracellular viral titres,^62^ we first tested the extent of viral entry at different transduction time points using an MOI of 0.02. The total viral RNA copies in transduced cells were quantified by qRT-PCR using the Lenti-X qRT-PCR titration kit every 24 hours post infection (h.p.i) for up to 72 h.p.i. Interestingly, we observed an exponential increase in the number of intracellular RNA copies with each additional day that the podocytes were exposed to the pseudoviral particle in culture (**Figure 2A**), confirming an increase in viral uptake with incubation time. Consistent with these results, western blot quantification of the relative amount of Gag-p24 (in the pseudovirus’ HIV backbone^63^) taken up by the podocytes each day post-infection (**Figure 2B**) corroborated the results of relative viral RNA quantification shown in **Figure 2A**. Furthermore, western blot analysis of protein lysates generated from uninfected (mock) podocytes, or podocytes infected with S-pseudotyped, bald pseudotyped and VSV-G pseudotyped virus for 72 hours revealed bands that correspond to the spike protein only in the lysate from podocytes infected with the S-pseudotyped virus. A band corresponding to β-actin (used as loading control) was present in all cell lysates while the band for p24 was observed in all the pseudotyped virus infected cell lysate but not the mock control, demonstrating the presence of the pseudoviral particle only in the infected cells (**Figure 2C**).

**Figure 2:**
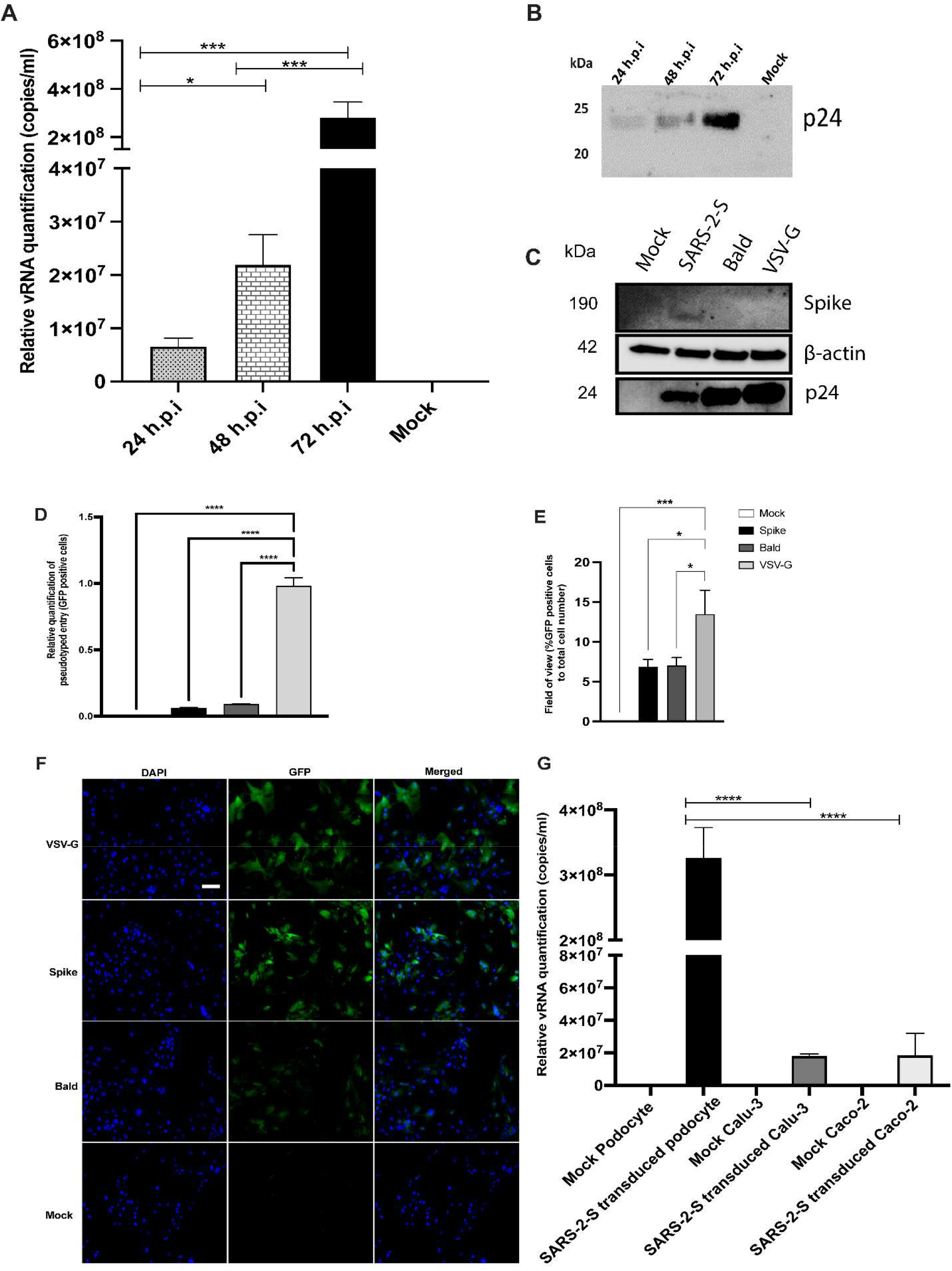
SARS-CoV-2 S-pseudovirus infection of human iPS cell-derived podocytes. (**A**) qRT-PCR analysis using Lenti-X titration kit revealed time-dependent increase in the copies of viral RNA in human iPS cell-derived podocytes infected with the S-pseudovirus. (**B**) Western blot confirmed time-dependent increase of S-pseudovirus particles in human iPS cell-derived podocytes, where p24 (GAG) is a marker for the pseudoviral capsid protein. (**C**) Western blot from cell lysates of mock and infected (S-, Bald and VSV-G pseudotyped) podocytes confirming the presence of SARS-CoV-2 spike proteins in S-pseudotyped infected podocytes, HIV viral protein p24 visible in all pseudotyped infected cells but not in mock and β-actin used as loading control present in all cell lysates (**D**) qRT-PCR data measuring the levels of EGFP using GFP specific primers (**E**) Percentage of EGFP positive cells compared to total cell number (**F**) Immunofluorescent staining showing DAPI, EGFP and merged in pseudotyped infected cells and mock. Scale bar: 100 µm. (**G**) qRT-PCR results showing significantly higher number of S-pseudotyped copies in human iPS cell-derived podocytes compared to Calu-3 and Caco-2 cell lines, 72 hours post infection (h.p.i.). The statistical test was done by One-way ANOVA with Sidak’s multiple comparison test. Error bars indicate standard deviation of the mean. Only p values of 0.05 or lower were considered statistically significant (p > 0.05 [ns, not significant], p < 0.05 [*], p < 0.01 [**], p < 0.001 [***], p < 0.0001 [****]).

After infection of the podocytes with the different pseudoviruses, the transcript level of GFP from infected and uninfected cells was quantified using qRT-PCR (**Figure 2D**). When compared to levels of viral uptake (corresponding tothe GFP mRNA levels) in VSV-G pseudotyped infected cells, there was a significantly lower uptake in cells infected with S-pseudotyped virus, which is expected since entry of VSV-G typed virus does not require specialized receptors as S-pseudotyped viruses do. GFP-positive cells were imaged by fluorescence microscopy and quantified relative to the total cell counts (shown by DAPI) (**Figure 2E**). The GFP expression results were comparable to that obtained with qRT-PCR with significantly more uptake in podocytes infected with VSV-G pseudotyped virus compared to the ones infected with S-pseudotyped virus. **Figure 2F** show representative microscopy images of infected and mock podocytes.

To examine how the levels of viral uptake in the podocytes compare to other organ-specific cell types, we examined pseudoviral uptake in Calu-3 (lung epithelial) and Caco-2 (colon epithelial) cells along with mock-infected cells (TBS control) for each cell type for 72 hrs. Intriguingly, there was significantly more viral uptake in the podocytes than Calu-3 (p-value < 0.0001) and Caco-2 cells (p-value < 0.0001) (**Figure 2G**). These results reveal high permissivity of human iPS cell-derived podocytes to S-pseudotyped viral infection.

### Live SARS-CoV-2 virus infects and replicates in human iPS cell-derived podocytes

To explore the susceptibility of human iPS cell-derived podocytes to live SARS-CoV-2, the podocytes were incubated with SARS-CoV-2 strain USA-WA1/2020 at MOI of 0.01, 0.1 or 1.0 for 1 hour. The range of MOIs was chosen based on previously established models of infection kinetics.^64^ After 1 hour incubation, cells were washed with PBS and then incubated with fresh culture medium for 24, 48 and 72 hours. Phase contrast images of the infected podocytes revealed changes in cell morphology at higher MOI compared to mock (**Supplementary Figure 1A)**. At 24, 48 and 72 hours post-infection, total RNA was extracted from both the cell pellets and the supernatant to evaluate both intracellular and extracellular viral RNA (vRNA) levels.

We quantified the intracellular and extracellular vRNA copies by qRT-PCR using primers specific for SARS-CoV-2 spike and nucleocapsid genes (normalized to endogenous control). Analysis of cell pellets collected at 24 and 48 h.p.i demonstrated high levels of viral RNA transcripts in cells infected with MOI of 1 (**Figures 3A and B**). At 72 h.p.i., higher levels of viral RNA transcripts were detected in the cells infected with MOI of 0.01 (**Figure 3C**). At 72 h.p.i, lower levels of viral RNA were detected in the intracellular fractions from the higher MOI of 0.1 and 1.0 conditions likey due to higher cellular toxicity at increasing MOI, leading to a lower number of healthy cells available for additional rounds of viral propagation. As expected, no viral RNA was detected in the mock-treated cells (**Figure 3C**). These results indicate high SARS-CoV-2 susceptibility and primary infection of the podocytes even at low MOI of 0.01. Quantification of the levels of spike and nucleocapsid in cell supernatants revealed an inverse trend (for 72 h.p.i) whereby significantly higher amounts of vRNA was detected in supernatants from cells infected with a MOI of 1.0 than in the cells infected with a MOI of 0.1 or 0.01 at 24, 48 and 72 h.p.i. (**Supplementary Figures 1B, C and D**).

**Figure 3:**
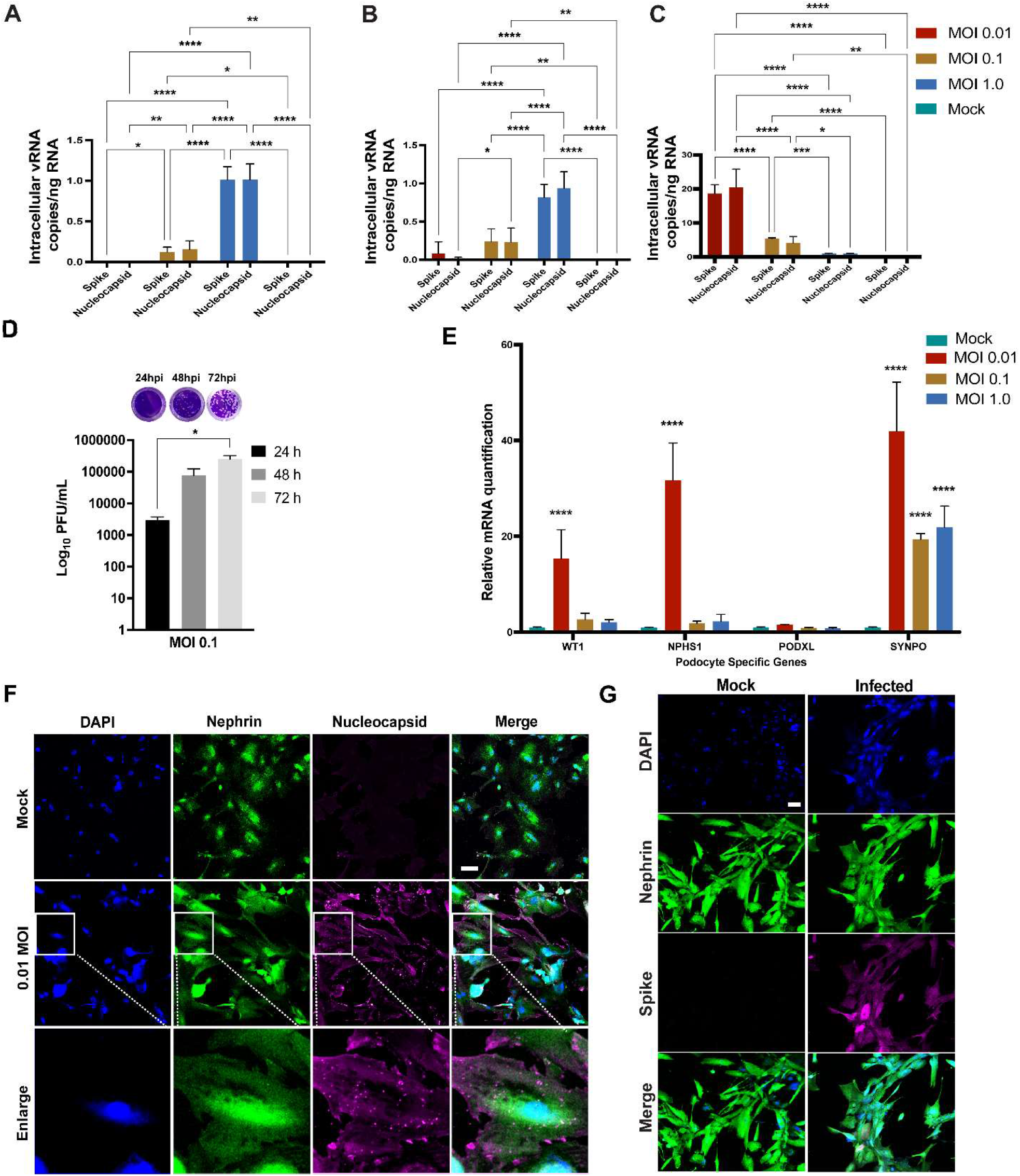
Susceptiblity of human iPS cell-derived podocytes to infection by live SARS-CoV-2. qPCR analysis of human iPS cell-derived podocytes infected with SARS-CoV-2 revealed intracellular uptake of the virus for 24 h.p.i (**A**), 48 h.p.i (**B**) and 72 h.p.i (**C**). (**D**) plaque assay quantification from supernatant obtained from infected podocytes at 24, 48 and 72 h.p.i. (**E**) qPCR analysis of podocyte-specific genes revealed that both synaptopodin (SYNPO) and podocalyxin (PODXL) are significantly upregulated after infection with SARS-CoV-2 at MOI of 0.01, whereas SYNPO is significantly upregulated at multiple MOIs, and PODXL shows no significant changes with viral infection. (**F**) Human iPS cell-derived podocytes treated with SARS-CoV-2 (at MOI of 0.01) immunostain positive for Nucleocapsid protein (magenta), indicating successful infection with the virus. The cells were immunostained also for the podocyte marker Nephrin (green) and counterstained with DAPI (blue). Scale bar: 100 µm (**G**) Spike positive cells Nephrin and DAPI as nuclear counterstain in the infected podocytes. Scale bar: 100 µm. One-way analysis of variance (ANOVA) with Sidak’s multiple comparison test was used to determine statistical significance. Only p values of 0.05 or lower were considered statistically significant (p > 0.05 [ns, not significant], p < 0.05 [*], p < 0.01 [**], p < 0.001 [***], p < 0.0001 [****]). Error bars indicate standard deviation of the mean.

We next performed plaque assays to measure the amount of infectious SARS-CoV-2 particles released from infected podocytes at 24, 48 and 72 h.p.i (**Supplementary Figures 1E, F and G**, respectively). Ten-fold dilutions of each cell supernatant were assed in duplicates as previously described.^64^ The number of plaque forming units (PFU) was significantly higher in the cells infected with MOI of 0.1 at 72 h.p.i compared to 24 and 48 h.p.i. (**Figure 3D**).). However, a lower number of PFU was observed in cells infected with MOI of 1 compared to cells infected with MOI of 0.1 at 72h.p.i. (**Supplementary Figures 1G**). These data suggest that the higher vRNA levels observed in Figure S1B, C, D accounts for vRNA released from dying cells that is not incorporated in new infectious particles. Taken together, these results confirm that SARS-CoV-2 can infect and replicate in human iPS cell-derived podocytes

To probe the level of cell death caused by infection with SARS-CoV-2 at 72 h.p.i, we compared the number of nuclei per field of view in the DAPI stained infected cells to that in the mock-infected control. We observed significantly (p-value < 0.0001) more cell death in the infected wells compared to the mock-infected controls (**Supplementary Figure 2A**). We then quantified the mRNA levels of apoptotic genes as well as necroptotic genes to examine whether SARS-CoV-2 can trigger both apoptosis and necroptosis (a form of cell death mediating secretion of inflammatory cytokines)^65^ in the infected podocytes. It was previously shown that SARS-CoV-2 can trigger apoptosis in Calu-3 cells through caspase-8 activation and that the process was dependent on viral replication.^66^ Similarly, our results showed a significant increase in Caspase 8 mRNA (p-value < 0.005) at MOI of 0.01, but not caspase 7, suggesting that the activation of cellular apoptosis is dependent on viral replication (**Supplementary Figure 2B**). To determine whether SARS-CoV-2 infected podocytes undergo necroptosis, we assessed mRNA expression of the mixed lineage kinase domain-like (MLKL) and the receptor-interacting protein kinase-3 (RIPK3), two effectors of necroptosis.^66, 67^ There was a significant upregulation of MLKL (p-value < 0.0001) and RIPK3 (p-value < 0.0014) in the MOI of 0.01 infected cells (**Supplementary Figure 2B**), where higher levels of intracellular vRNA were detected (**Figure 3C**). These results are consistent with a prior report using Calu-3 cells, where activation of necroptosis pathway was shown to be dependent on viral replication.^66^ Conversely, no upregulation of MLKL or RIPK3 mRNA was observed in podocytes infected with either 0.1 or 1.0 MOI of SARS-CoV-2, **(Supplementary Figure 2B**), presumably due to the lower levels of viral replication in those conditions (**Figure 3C**). These data suggest that the podocytes are susceptible to cell death through the activation of necroptosis and apoptosis pathways when infected by SARS-CoV-2.

### SARS-CoV-2 infection alters podocyte-specific gene expression

We next examined whether SARS-CoV-2 infection affects the expression of genes involved in podocyte lineage specification. Changes in the expression levels of podocyte-specific genes and proteins often correlate with the onset and progression of podocytopathies,^68–70^ including HIVAN.^71^ Additionally, defects in podocyte structure and function to their detachment from the glomerular basement membrane and subsequent loss of the cells into urine, and the onset of glomerulopathies.^72–74^

Quantification of podocyte lineage determination genes (WT1, nephrin, podocalyxin and synaptopodin) after SARS-CoV-2 viral infection revealed significant increase in the expression of WT1, NPHS1 and SYNPO at MOI of 0.01 compared to mock, and a marginal increase in the expression of PODXL (**Figure 3E and Supplementary Table 1**). The increased expression of nephrin may result from compensatory mechanism to help maintain podocyte physiology post-infection and minimize destabilization of their cellular phenotype as previously reported in a diabetic model of podocyte injury ^75^. The increase in nephrin gene expression also correlated with the presence of more foot-like processes in the podocytes infected with SARS-CoV-2 at an MOI of 0.01 (**Supplementary Figure 2C**). However, at MOI of 1.0, we observed changes reminiscent of foot process retraction with a concomitant reduction in nephrin mRNA expression (**Figure 3E & Supplementary Figure 2C**) indicating a possible maladaptive response with increased viral infection burden. These results indicate that SARS-CoV-2 infection of human iPS cell-derived podocytes leads to dynamic changes in the expression of podocyte-specific genes. Together, our results suggest that infection of podocytes by SARS-CoV-2 results in disrupted molecular profile as well as structural changes which can lead to cell detachment and death (**Supplementary Figure 2A**).

Immunofluorescence analysis of the SARS-CoV-2 infected human iPS cell-derived podocytes showed positive immunostaining of the nucleocapsid protein at MOI of 0.01, revealing the presence of viral proteins with in the cytoplasm even at low MOI (**Figure 3F**). This result further confirmed our observation that SARS-CoV-2 can establish active infection in human iPS cell-derived podocytes. The infected podocytes also exhibited plaque-like structures (**Supplementary Figure 2D**) indicating loss of cell membrane integrity. Coronaviruses have been shown to manipulate cell cycle progression by hijacking the host DDXi RNA helicases to enhance their replication^5^. Furthermore, syncytia formation has been reported in SARS-CoV-2 infected cells^76, 77^. We observed a more pronounced DAPI staining and spreading indicating more nuclear content in SARS-CoV-2 infected podocytes compared to the mock samples (**Figure 3F, 3G and Supplementary Figure 2E**). Changes in cellular nuclear content correlates with enhanced viral replication,^5^ and this could potentially result from SARS-CoV-2 infection of the podocytes producing syncytia-like phenotype that could be explored futher in future studies. Additionally, podocytes infected with SARS-CoV-2 were immunoreactive to anti-spike antibody, which further suggest the presence of an active viral infection (**Figure 3G**).

### Human iPS cell-derived podocytes express several spike-interacting factors

Previous studies have identified specific host factors that can facilitate entry of SARS-CoV-2 virus into various tissues and cell types,^18, 19, 78, 79^ but it is unclear which viral uptake and processing factors are expressed in fully differentiated human iPS cell-derived podocytes. To examine this, we first used BioGRID (a database of protein, genetic, and chemical interactions) to identify proteins that have been experimentally shown to have spike-interacting capabilities. This analysis provided twenty-four factors involved in the binding and processing of SARS-CoV-2 through the spike protein (**Supplementary Figure 3A & Supplementary Table 2**). We then examined the gene expression levels of the twenty-four spike-interacting factors in human iPS cell-derived podocytes using our previously generated microarray data (**Figure 4A**).^54^ Intriguingly, the podocytes expressed twenty (out of twenty-four) spike interacting factors. Among them, BSG/CD147, ACTR3, PLS3, ZDHHC5, and GOLGA7 were highly expressed in the human iPS cell-derived podocytes (**Figure 4A & Supplementary Table 2**). These results indicate that human iPS cells possess many of the factors involved in SARS-CoV-2 binding and processing, which further supports our data from above showing high SARS-CoV-2 infectivity in the podocytes.

**Figure 4:**
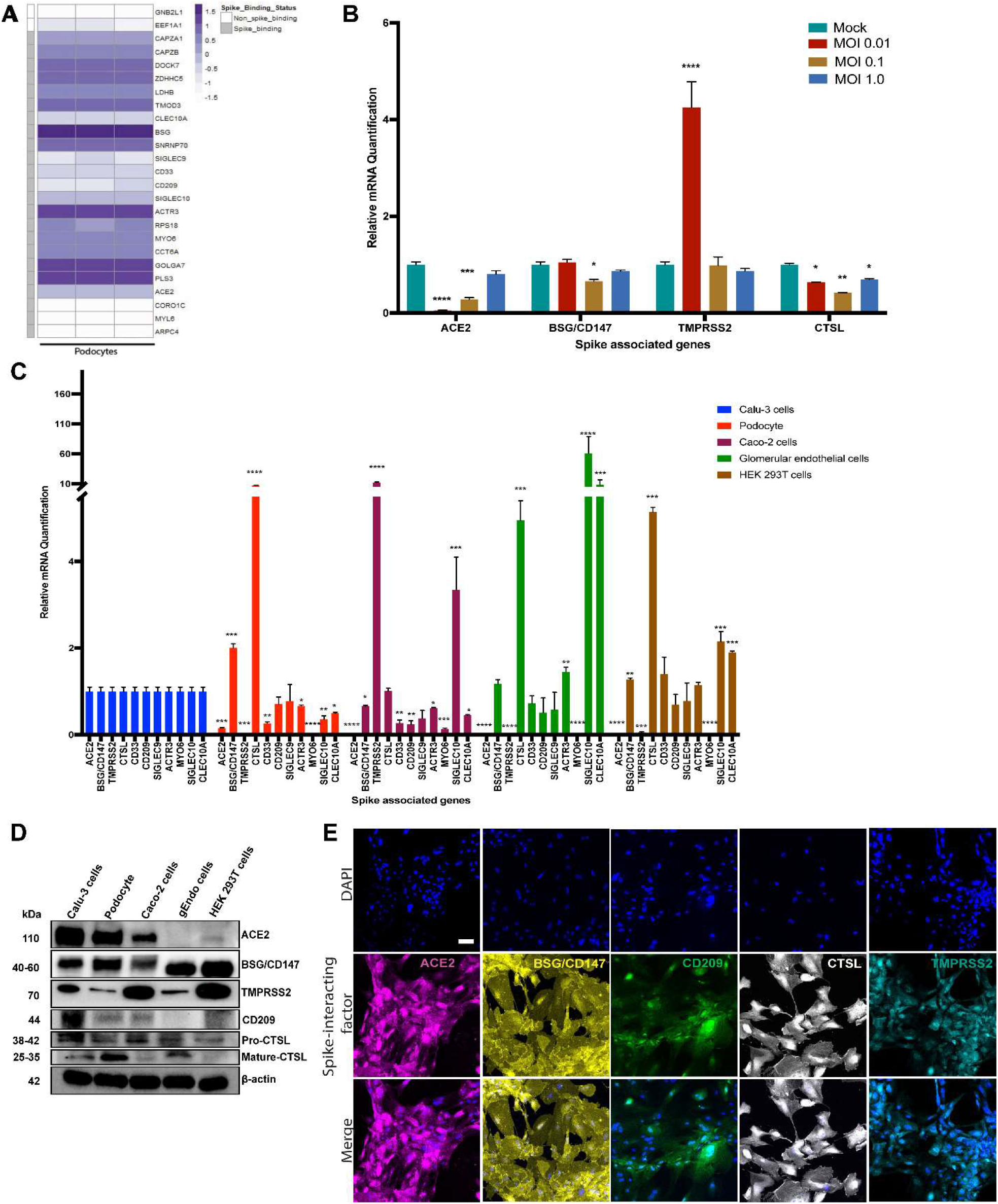
Transcriptomic and protein level analyses of spike-interacting factors in mature podocytes differentiated from human iPS cells. (**A**) Heatmap showing expression levels of spike associated genes from three biological human iPS cell-derived podocyte samples. SIGLEC9, Sialic acid-binding Ig-like lectin 9; CAPZA1, F-actin-capping protein subunit alpha-1; CLEC10A, C-type lectin domain family 10 member A; CD33, Myeloid cell surface antigen CD33; TMOD3, Tropomodulin-3; ACE2, Angiotensin Converting Enzyme 2; BSG/CD147, Basigin/CD147 molecule; CD209, CD209 Antigen; MYO6, Unconventional myosin-VI; PLS3, Plastin-3; LDHB, L-lactate dehydrogenase B chain; GNB2L1/RACK, Receptor of activated protein C kinase 1; SNRNP70, U1 small nuclear ribonucleoprotein 70 kDa; DOCK7, Dedicator of cytokinesis protein 7; RPS18, 40S ribosomal protein S18; CAPZB, F-actin-capping protein subunit beta; GOLGA7, Golgin subfamily A member 7; ZDHHC5, Palmitoyltransferase ZDHHC5; SIGLEC10, Sialic acid-binding Ig-like lectin 10; ACTR3, Actin-related protein 3; MYL6, Myosin light polypeptide 6; CORO1C, Coronin-1C; ARPC4, Actin-related protein 2/3 complex subunit 4; CCT6A, T-complex protein 1 subunit zeta; (**B**) The expression of Spike associated genes (ACE2, BSG/CD147) and Spike processing genes (TMPRSS2, CTSL) are significantly impacted by infection at MOIs of 0.01 and 0.1, respectively. (**C**) qPCR quantification of nine of the human spike associated gene from (A) including Transmembrane Serine Protease 2 (TMPRSS2) or cathepsin L (CTSL) in human iPS cell-derived podocytes, Calu-3, Caco-2, glomerular endothelial cells, and HEK 293T cells (normalized to Calu-3 groups). (**D**) Western blot analysis to evaluate protein expression of ACE2, BSG/CD147, CD209, TMPRSS2 and CTSL in the different cell type used; Calu-3, human iPS cell-derived podocytes, Caco-2, glomerular endothelial (gEndos)cells, and HEK 293T cells. (**E**) Immunocytochemistry analysis of ACE2, BSG/CD147, CD209, TMPRSS2 and CTSL expression in iPS cell-derived podocytes. Scale bar: 100 µm. One-way analysis of variance (ANOVA) with Sidak’s multiple comparison test was used to determine statistical significance. Only p values of 0.05 or lower were considered statistically significant (p > 0.05 [ns, not significant], p < 0.05 [*], p < 0.01 [**], p < 0.001 [***], p < 0.0001 [****]). Error bars indicate standard deviation of the mean.

Because BSG/CD147 was the most highly expressed gene among the spike-interacting factors expressed in podocytes (**Figure 4A**), we wondered if this receptor might be employed by the SARS-CoV-2 virus to directly infect the cells. We initially examined the expression levels of BSG/CD147 in SARS-CoV-2-infected podocytes. We also examined the expression of ACE2 given its involvement in SARS-CoV-2 binding and infection of many cell types,^11, 16, 63, 80^ as well as cell surface protease TMPRSS2^20^ and endosomal Cathepsin L (CTSL). We quantified the relative expression of these genes in podocytes infected with SARS-CoV-2 for 72 h using different MOIs (**Figure 4B**). We observed that infection at MOI of 0.01 and 0.1 lead to significant reduction in ACE2 expression when compared to uninfected podocytes. Compared to the mock condition, the expression levels of BSG/CD147 remained unchanged for MOI of 0.01 and 1.0, but decreased significantly when the podocytes were infected at an MOI of 0.1. In addition, TMPRSS2 expression was significantly increased with SARS-CoV-2 infection at MOI of 0.01 but remained relatively similar to the mock condition when the cells were infected at MOI of 0.1 and 1.0. CTSL expression was significantly reduced in all three MOIs. These results show that at MOI of 0.01, expression of ACE2 decreases but that of TMPRSS2 increases, suggesting enhanced enzymatic activity necessary to cleave Spike for processing. These results also show that the low MOI of 0.01 is sufficient for the infection of human iPS cell-derived podocytes with SARS-CoV-2 and indicate that SARS-CoV-2 infection of the podocytes leads to dynamic changes in the expression of spike-binding factors (**Figure 4B**) as well as podocyte-specific genes (**Figure 3C**).

### Comparative analysis of spike-associated genes in podocytes and other tissue-specific cells

ACE2 is expressed in a variety of human tissues and has been shown to function by counter-balancing the renin-angiotensin-aldosterone system.^80, 81^ We quantified the basal mRNA expression levels of ACE2, BSG/CD147 and other spike-associated genes in different human cell types (**Figure 4C**). The expression of these genes is important for the uptake and cleavage of the spike glycoprotein, fusion of SARS-CoV-2 and cell membranes, and subsequent release of viral genome into the cytoplasm of an infected cell.^60^

Studies have shown that some cells with little to no ACE2 mRNA or protein expression can still be infected with SARS-CoV-2,^15, 81^ suggesting that other class of receptors might facilitate viral infection in ACE2-deficient cell types. Additionally, it has been shown that the expression levels of viral uptake receptors can vary significantly between different cell types.^18, 78^ These findings suggest that ACE2 may not be the primary or the only receptor for SARS-CoV-2 in some cells, and that there could be multiple mechanisms for viral infection and processing. For example, BSG/CD147 has been shown to mediate viral uptake in Vero and HEK 293T cells^18, 19^ but was suggested to not be directly involved in SARS-CoV-2 infection of lung cells (e.g Calu-3).^82^ Our BioGrid results indicated that BSG/CD147 is one of the most highly expressed spike-interacting factors in the human iPS cell-derived podocytes. We further examined expression levels of several of the factors in multiple cell types (podocytes, Calu-3, Caco-2, glomerular endothelium, and HEK 293T) to help understand the levels of tissue or cell-type specificity (**Figure 4C**). We found that there was no ACE2 expression in HEK 293T cells, consistent with a previous report.^24^ We also found little to no ACE2 expression in glomerular endothelial cells and Caco-2 cells. Intriguingly, ACE2 expression in human iPS cell-derived podocytes is approximately 10 times lower than the expression level in Calu-3 cells and slightly higher than the expression level in Caco-2 cells. This result was surprising as our results showed that podocytes were much more permissive to infection by SARS-CoV-2 pseudovirus compared to Calu-3 and Caco-2 cells (**Figure 2G**).

SARS-CoV-2 entry into cells can also be facilitated by the transmembrane protease TMPRRS2. We observed significantly low expression levels of TMPRRS2 in the podocytes compared to Calu-3 cells (**Figure 4C**). A lower level expression of TMPRRS2 was also previously reported for cardiomyocytes derived from human embryonic stem cell (hESC-CMs) and a different endosomal viral processing protease was shown to be much more highly expressed.^24^ Our qPCR results showed a significantly higher expression of CTSL in podocytes when compared to Calu-3 cells (Figure 4B), suggesting that the mechanism of SARS-CoV-2 entry in hESC-CMs^24^ and in podocytes might be different from the TMPRRS2-dependent mechanisms observed in lung epithelial cells.^16, 63^

It is likely that SARS-CoV-2 infection of podocytes relies on ACE2, BSG/CD147 and other genes that might direct membrane fusion and/or entry through the endo-lysosomal pathway. Together, these results show that human iPS cell-derived podocytes express proteins that make them susceptible to SARS-CoV-2 infection, similar to human iPS cell derived cardiomyocytes.^24, 25^ The expression of CD209, which is recognized as an alternative receptor for lung and kidney epithelial and endothelial cells,^79^ was comparable between Calu-3 cells, human iPS cell-derived podocytes, glomerular endothelial cells and HEK 293T cells but significantly lower in Caco-2 cells (**Figure 4C**). The mRNA expression of the other genes, SIGLEC9, ACTR3, MYO6, SIGLEC10 and CLEC10A, varied between the different cell types (**Figure 4C**). Based on all these findings, we hypothesized that human kidney podocytes might employ other or multiple spike-binding receptors (not just ACE2) for SARS-CoV-2 viral uptake. We examined this possibility through antibody blocking experiments as described below.

#### Podocytes express spike associated factors at the protein level

We validated the relative expression of three uptake (ACE2, BSG/CD147, CD209) and two processing (TMPRSS2, and CTSL) factors at the transcriptome and proteomic levels in podocytes and other cell types (Calu-3, Caco-2, glomerular endothelia and 293T cells) using qRT-PCR, western blot, and immunocytochemistry analyses. Transcript levels of ACE2 was highest in Calu-3 cells followed by podocytes and then Caco-2 (**Figure 4C**). This was confirmed in the western blot analysis that revealed higher expression of ACE2 in Calu-3 cells and then podocytes with no expression in glomerular endothelial cells (gEndos) (**Figure 4D**). This result also demonstrated that podocytes express more ACE2 than Caco-2 cells, consistent with the gene expression data in figure 4C. Additionally, the western blot data for BSG/CD147 corroborated the mRNA data showing higher expression in podocytes than Calu-3 and Caco-2. Figure 4D also shows the expression of CD209 in podocytes and the other cell types. We observed relatively low gene and protein level expression of TMPRSS2 in podocytes when compared to the other cell types. Although mature CTSL protein is present in only podocytes, Calu-3 and glomerular endothelia cells, pro-CTSL is present in all the cell types which may explain the presence of the CTSL mRNA in all the cell types even when they do not express mature CTSL protein. Finally, expression of all these proteins was also validated using immunocytochemistry in podocytes (**Figure 4E**), Calu-3 (**Supplementary Figure 3B)**, Caco-2 (**Supplementary Figure 3C)**, glomerular endothelial (**Supplementary Figure 3D)** and 293T (**Supplementary Figure 3E)** cells.

### Receptor antibodies can reduce SARS-CoV-2 pseudovirus entry into human iPS cell-derived podocytes

It is known that coronaviruses can enter cells via direct fusion at the cell surface or internalization through the endosomal compartment ^60, 83^. Based on the hypothesis that SARS-CoV-2 could exploit both ACE2 and BSG/CD147 receptors for viral uptake in human iPS cell-derived podocytes, we investigated whether antibodies against these two spike receptors can block the entry of S-pseudotyped virus in the cells. We used anti-hACE2 and anti-CD147 (anti-BSG) antibodies at varying concentrations to block ACE2 and BSG/CD147 receptors from interacting with pseudoviral particles. An S-pseudotyped virus MOI of 0.02 was used for all antibody blocking experiments. Podocytes were lysed after 60 h.p.i and quantified for viral uptake by measuring the reporter gene using qPCR analysis with data normalization to GAPDH using the delta-delta Ct (2^—ΔΔ^*Ct*) method.

When the podocytes were blocked with ACE2 or BSG/CD147 antibody at varying concentrations (from 0.1 to 5 µg/ml), we observed a concentration-dependent and statistically significant decrease in viral uptake (**Figure 5A**). The highest concentration of the antibody (5 µg/ml) was most effective for blocking the receptors while significantly (p-value < 0.0001) reducing cellular uptake of the virus (**Figure 5A**). These results confirm that ACE2 and BSG/CD147 facilitate SARS-CoV-2 S-pseudotyped viral update in podocytes. Thus, the observed high expression of BSG/CD147 receptor in podocytes revealed by microarray (**Figure 4A**), qPCR data (**Figure 4C**), western blotting (**Figure 4D** and immunocytochemistry (**Figure 4E**) further suggest that these receptors interact with the spike protein of SARS-CoV-2 and facilitate its uptake and entry into the cells. As a result, blocking with anti-BSG/CD147 significantly decreased viral uptake similar to that observed with ACE2 blocking.^16, 25^

**Figure 5:**
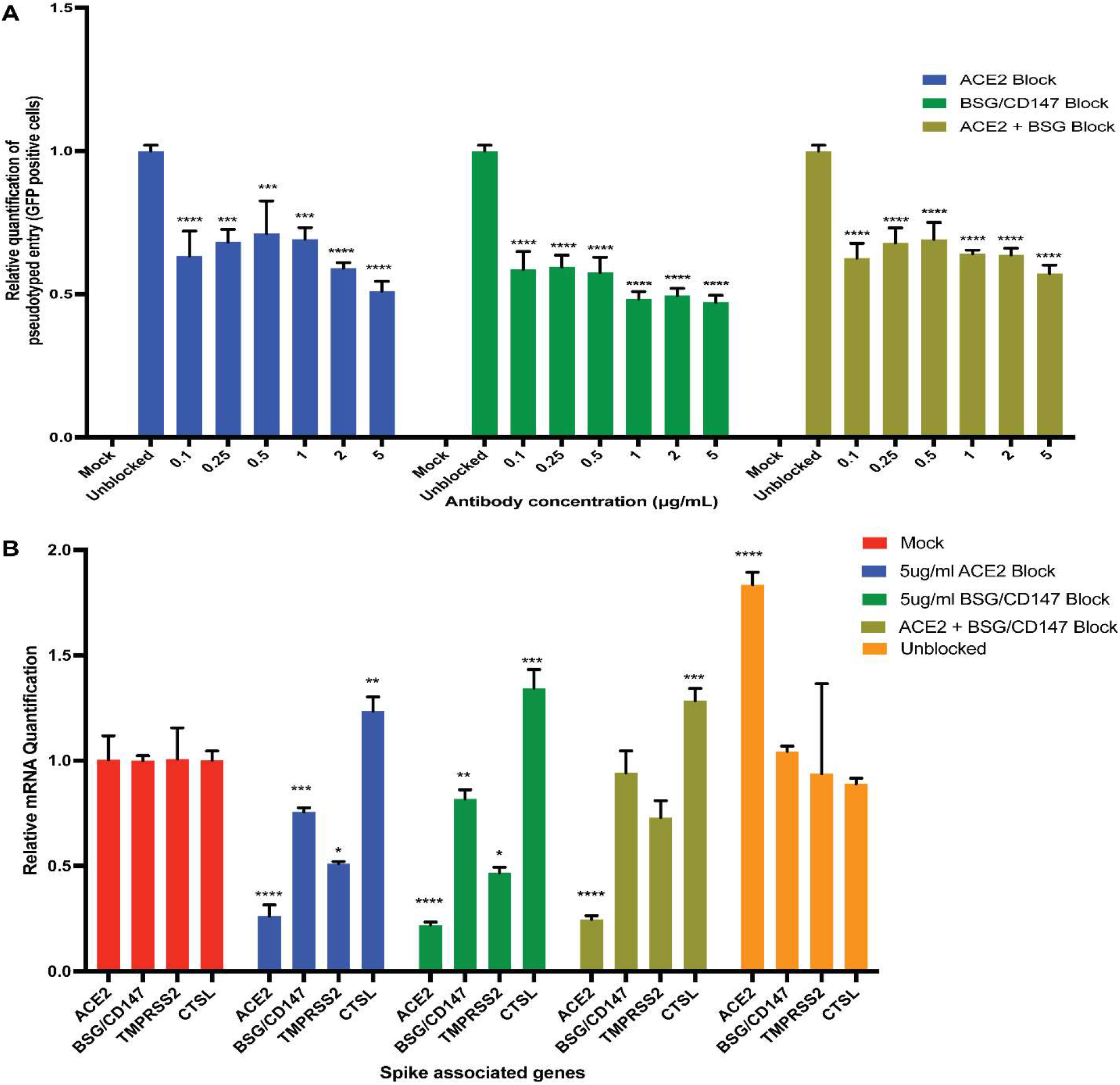
Antibody blocking reveal roles of ACE2 and BSG/CD147 receptors in viral uptake. (**A**) qPCR quantification of S-pseudotyped entry relative to antibody concentration (normalized to unblocked samples). Human iPS cell-derived podocytes were incubated with different dilutions of anti-ACE2, anti-BSG or both for an hour followed by infection with S-pseudotyped virus at MOI 0.02 for 60 h. (**B**) qPCR quantification of Spike binding receptor genes (ACE2, BSG/CD147) and Spike processing factor genes (TMPRSS2, CTSL) at 5 µg/ml for both anti-ACE2 antibody blockage and anti-BSG antibody blockage showing significant changes in gene expression with optimal receptor blockage when compared to unblocked samples (normalized to mock groups). One-way analysis of variance (ANOVA) with Sidak’s multiple comparison test was used to determine statistical significance. Only p values of 0.05 or lower were considered statistically significant (p > 0.05 [ns, not significant], p < 0.05 [*], p < 0.01 [**], p < 0.001 [***], p < 0.0001 [****]). Error bars indicate standard deviation of the mean.

To examine if blocking with both anti-ACE2 and anti-BSG had a synergistic effect, the podocytes were simultaneously pretreated with both the antibodies and then infected with the virus. We observed similar trend where the lowest viral uptake was at 5 µg/ml (**Figure 5A**). Pre-treatment of infected podocytes with both anti-ACE2 and anti-BSG/CD147 antibodies at a concentration as low as 0.1 µg/ml significantly diminished viral uptake (p-value < 0.0001), suggesting that both ACE2 and BSG/CD147 are involved in SARS-CoV-2 internalization in human iPS cell-derived podocytes.

Next, we examined whether the expression of the key viral entry genes, ACE2, BSG/CD147, TMPRSS2 and CTSL are altered during antibody blocking. Exposure of podocytes to the pseudoviral particles after receptor blocking with 5 µg/ml antibody concentrations (for both anti-ACE2 and anti-CD147 (anti-BSG)) resulted in changes in the expression of spike-associated genes when compared to the unblocked control samples (**Figure 5B**). The genes showed diminished expression when blocked with an antibody, which could be attributed to blockage of the cell surface receptor and prevention of viral entry. This result suggests that although ACE2 might be the primary SARS-CoV-2 entry receptor in other tissues, podocytes utilize both ACE2 and BSG/CD147 for efficient viral binding and uptake.

## Discussion

The global pandemic caused by SARS-CoV-2 has resulted in the loss of millions of lives and caused devastating social and economic burdens. The disease mostly presents as a respiratory illness, similar to viral pneumonia, and in more severe cases as acute respiratory distress syndrome (ARDS).^4, 5, 21^ In addition, clinical and *in vitro* studies have revealed the multi-organ effects of the disease,^27, 32^ in which several COVID-19 patients simultaneously experience renal, cardiac, neurological, digestive, and/or pancreatic complications.^15, 21^ SARS-CoV-2 RNA has been detected in the kidney glomerulus of patients who have died from the disease.^29^ Although the onset of acute kidney injury and collapsing glomerulopathy^34^ have been clinically associated with the severe form of COVID-19,^32^ it remains unknown how the kidneys are specifically targeted by the virus, and whether podocytes – the specialized epithelial cells that help form the blood filtration barrier in the kidneys – can be directly targeted by the virus. Technologies that can help understand these processes in human cells and tissues could advance current understanding of the disease and enable therapeutic discovery.

To help address some of these important questions, we show in this report that human iPS cell-derived podocytes, which have similar genotypic and phenotypic characteristics of the mature and specialed human kidney podocytes (Musah et al., 2017) are highly susceptible to SARS-CoV-2 infection even at low MOI of 0.01 (**Figure 3A, B and C**). Initial viral infection of cells at low MOIs (< 1.0) allows additional rounds of virus replication in in a 2D culture, as the replicating virus can enter adjacent cells that were not infected during primary infection. Indeed, we observed higher levels of intracellular vRNA and higher number of PFU at 72 h.p.i. in cells infected with MOI of 0.01 of SARS-CoV-2 compared to MOI of 1.

These results are relevant and consistent with a recent report showing that SARS-CoV-2 isolated from COVID-19 autopsied kidney could infect kidney tubular cells *in vitro* and lead to extensive viral replication that produced 1000-times increased in viral RNA, confirming the presence of infectious virus in the kidneys.^32^

Due to the lower ACE2 mRNA expression in podocytes when compared to Calu-3, we evaluated the extent of viral infections using various MOIs from 0.01 to 1.0, as it was initially unknown if human iPS cell-derived podocytes would be permissive to direct infection with SARS-CoV-2. Typically, high MOIs (e.g., 1.0 or more) are required if the cell type is minimally permissive to viral infection^4^ as observed for some organoid models^49, 84^ within which cell-type-specific responses could not be fully evaluated due to high levels of heterogeneity.^85^ We show, however, that specialized podocytes derived from human iPS cells can be directly infected with SARS-CoV-2 at low MOIs of 0.01 to 1 (**Figure 3C**). The level of the cellular uptake of viral particles can be quantified using multiple assays including immunofluorescence microscopy (for the structural protein N or S, or against dsRNA intermediate), by quantitative RT-PCR of vRNA and plaque assay to quantify infection in the supernatant. ^4^ Indeed, we confirmed viral uptake by the human iPS cell-derived podocytes using qRT-PCR at 24, 48 and 72 h.p.i. (Figures 3A-C), plaque assay (Figure 3D) and immunofluorescence microscopy analysis with both anti-N and -S antibodies (**Figures 3F and G)**.. In addition, our investigation of how infection alters the expression of the spike-associated genes revealed a significant reduction in ACE2 expression when compared to uninfected samples (**Figure 4B**). This is in line with down regulation of ACE2 expression upon SARS-CoV spike protein binding which promotes lung injury^86^ as well as reduction in ACE2 expression due to SARS-CoV replication in Vero cells.^87^

Our results also revealed that human iPS cell-derived podocytes express lower levels of ACE2^88^ and TMPRSS2 when compared to Calu-3 (**Figure 4C, D and E**). Since SARS-CoV-2 - host interaction is vital for viral pathogenesis, ultimately determining the outcome of infection,^5^ and the functional activity of the virus depends on the proteolytic processing during cell entry,^83, 89^ we next sought to identify other factors that could mediate viral entry in iPS cell-derived podocytes. We utilized BioGRID analysis to gain insight into SARS-CoV-2 – host interactions by mapping out spike-binding proteins expressed in podocytes. Viral processing factors have been shown to be co-expressed with the type of spike binding receptor used by a given cell.^7, 20, 24^ In this study, we identified BSG/CD147 as a mediator of viral entry in podocytes, and CTSL as its putative processing factor (**Figure 4C and D**). We used qPCR, western blotting and immunocytochemistry to quantify the level of expression of different receptors and processing factors. We observed that both BSG/CD147 and CTSL are expressed at high levels in iPS cell-derived podocytes (**Figure 4C, D and E**), leading us to believe that they would be sufficient for SARS-CoV-2 entry, similar to what was reported in iPS cell-derived cardiomyocytes.^90^

The importance of employing cell models with mature phenotypes, which has historically been difficult for organoids and other iPS derived cell models, cannot be over-emphasized. For example iPS cell-derived renal organoids generate glomeruli with transcriptomic signatures similar to fetal stages^51^ which poses a question as to whether human iPS cell-derived cells can recapitulate the biology of SARS-CoV-2 infection in adults since vertical infection of the fetus is still unclear^91^ but remains a possibility^7, 92^. Furthermore, this points to the tissue-specific viral tropisms that may determine whether a productive infection is established in any given tissue. Therefore, it is important to understand these non-canonical SARS-CoV-2 entry-mediating proteins (i.e., other than ACE2 and TMPRSS2) so that we can establish effective methods to block viral replication in those tissues in which ACE2/TMPRSS are poorly expressed or not employed for viral infection.

Aside from being a receptor for SARS-CoV-2, ACE2 plays important role in different tissues in controlling blood pressure^18, 20, 93^ or preventing heart failure and kidney injury. ^88, 94, 95^ As such, development of drugs to block ACE2 might have a negative effect on its other protective functions. BSG/CD147 or EMMPRIN is a transmembrane glycoprotein belonging to the immunoglobulin superfamily.^17^ BSG/CD147 has been implicated in tumor metastasis, inflammation and viral infection^17, 96–98^ and also previously shown to facilitate SARS-CoV invasion in host cells.^17, 18^ Our results shows that antibody blocking of BSG/CD147 receptors significantly reduces SARS-CoV-2 viral uptake in human iPS cell-derived podocytes (**Figure 5A**). Thus, BSG/CD147 could potentially be a useful target for antiviral therapeutics including those aimed to address SARS-CoV-2 infections and COVID-19 disease.

## Author Contributions

T.D.K. and S.M. conceived the strategy for this study, designed the experiments, interpreted the results, and wrote the manuscript. T.D.K performed the experiments with assistance from co-authors. T.D.K. and R.B differentiated podocytes from human iPS cells; T.D.K. and M.B. designed BSL3-level experiments for SARS-CoV-2 viral infection of cells, M.B. performed the infection experiments, and T.D.K. and M.B. analyzed the data; T.T. performed viral RNA isolation from podocytes infected with live virus; R.B. created the schematic illustrations. X.M, helped with confocal microscopy; M.A.B. and T.D.K analysed microscopy data. A.E.O. performed the BioGRID analysis and generated heatmaps from microarray data previously generated by S.M.; All authors contributed to data analyses, figure preparations, discussion of results and the writing and editing of the manuscript.

## Acknowledgements

The authors thank all the members of Musah lab at Duke University for comments on the manuscript, and Dr. X. Campilongo for helpful comments on the manuscript.

## Data and code availability

The microarray data used in this study had been previously published.^54^

