## Supplementary Figures for "BSG/CD147 and ACE2 receptors facilitate SARS-CoV-2 infection of human iPS cell-derived kidney podocytes"

### Supplementary Figures and Legends

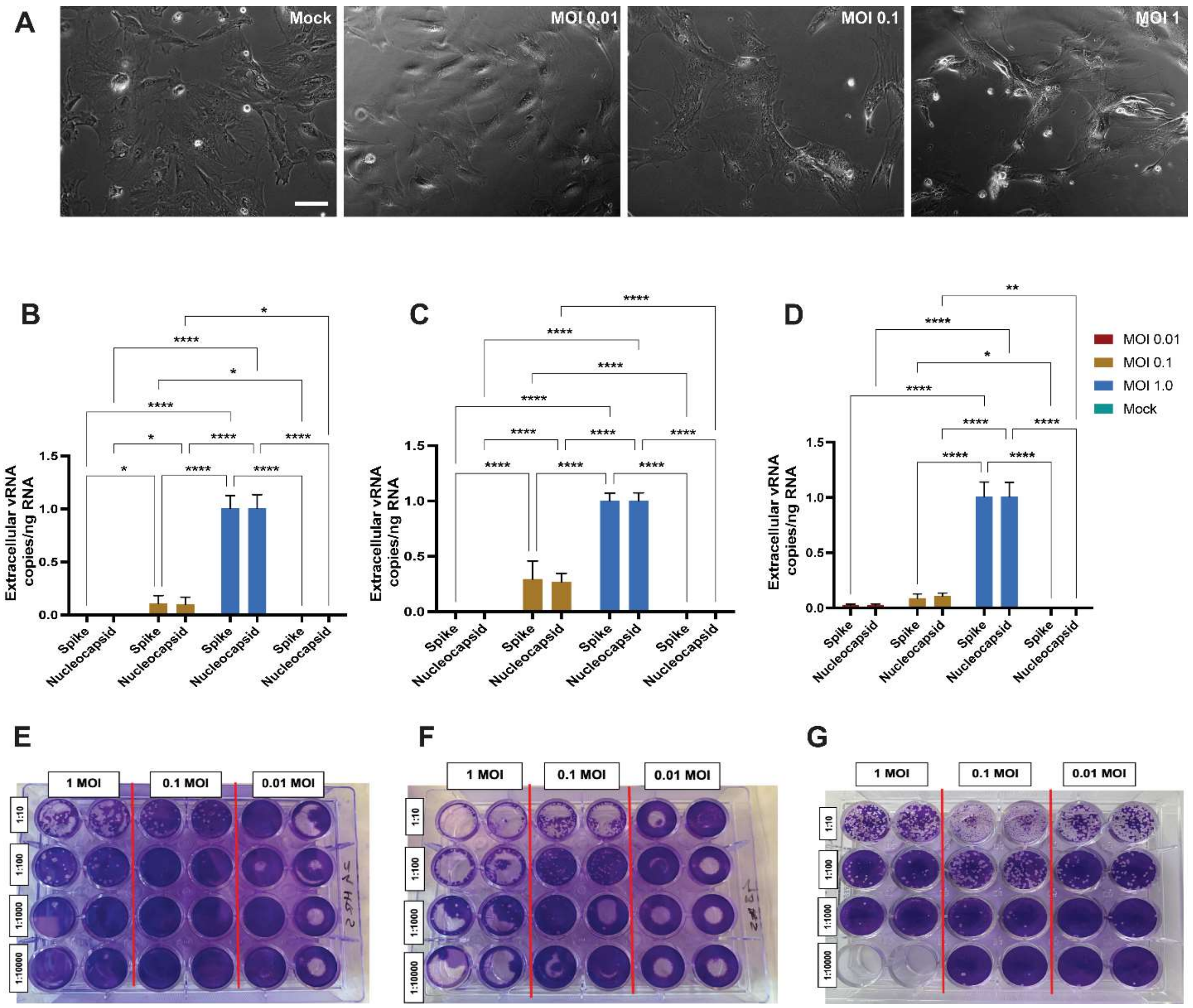


**Supplementary Figure 1: Effect of SARS-CoV-2 infection on human iPS cell-derived podocyte on cell viability.** (A) Phase contrast images of human iPS cell-derived podocyte morphology after 48 h.p.i at MOI of 0.01, 0.1 and 1 revealing changes in morphology when compared to the mock. Scale bar: 100 µm (**B-D**) qPCR analysis of human iPS cell-derived podocytes infected with SARS-CoV-2 confirmed release of the virus into the media (extracellular) for 24 h.p.i (**B**), 48 h.p.i (**C**) and 72 h.p.i (**D**). (**E-G**) Plate showing presence of plaques for the plaque assay quantification from supernatant obtained from infected podocytes at (**E**) 24 h.p.i, (**F**) 48 h.p.i and (**G**) 72 h.p.i.

The statistical test in this section was done by One-way ANOVA with Sidak's multiple comparison test. Error bars indicate standard deviation of mean. Only p values of 0.05 or lower were considered statistically significant ($p > 0.05$ [ns, not significant], $p < 0.05$

[*], p < 0.01 [**], p < 0.001 [***], p < 0.0001 [****]). For all statistical analyses, the GraphPad Prism 9 software package was used (GraphPad Software).

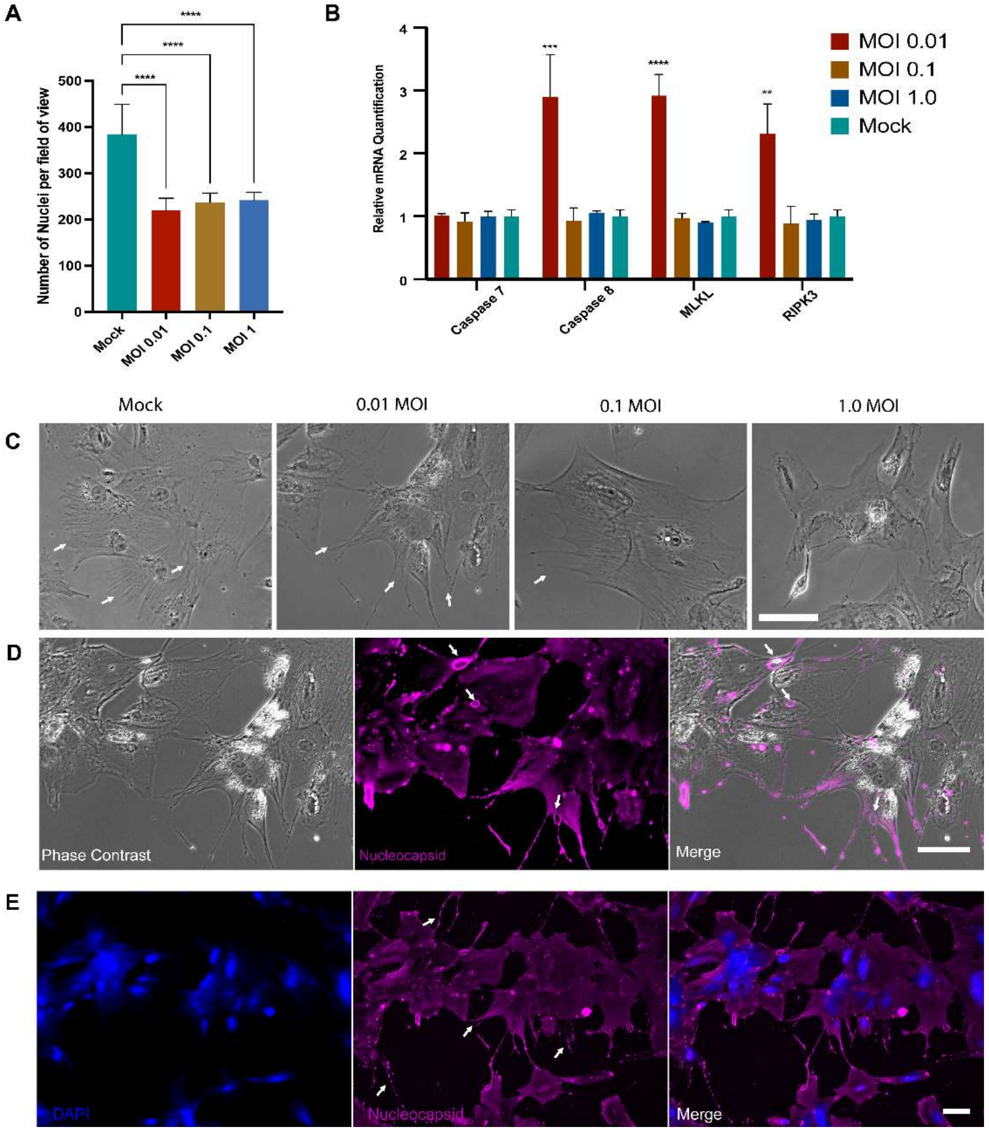


**Supplementary Figure 2**: Infection of podocytes result in phenotypic changes to the cells **(A)** DAPI count – number of nuclei per field of view. The number of nuclei in 5 field of view in each infection condition was counted using Image J and compared with the mock (**B**) qPCR analysis of necroptosis and apoptotic genes reveal significant increase in these genes which is dependent on viral replication. MLKL - mixed lineage kinase domain-like, RIPK3 - receptor-interacting protein kinase-3. **(C)** Phase contrast images of Mock and infected podocytes at MOI of 0.01, 0.1 and 1.0 indicating progression of loss of foot processes (white arrows). Mock and MOI of 0.01 cells possess foot processes while there is reduction in foot processes in MOI of 0.1 and there is no visible foot process projection in MOI of 1.0. **(D)** Phase contrast images and nucleocapsid staining of human iPS cell-derived podocytes treated with SARS-CoV-2 (MOI of 0.01) showing putative plaque formation (white arrows). **(E)** Human iPS cell-derived podocytes treated with SARS-CoV-2 (MOI of 0.01) stain positive for Nucleocapsid protein with enhanced foot processes (white arrows) with pronounced DAPI staining and spreading to the cell body. Scale bar: 100 µm

The statistical test in this section was done by One-way ANOVA with Sidak’s multiple comparison test. Error bars indicate standard deviation of mean. Only p values of 0.05 or lower were considered statistically significant ($p > 0.05$ [ns, not significant], $p < 0.05$ [*], $p < 0.01$ [**], $p < 0.001$ [***], $p < 0.0001$ [****]). For all statistical analyses, the GraphPad Prism 9 software package was used (GraphPad Software).

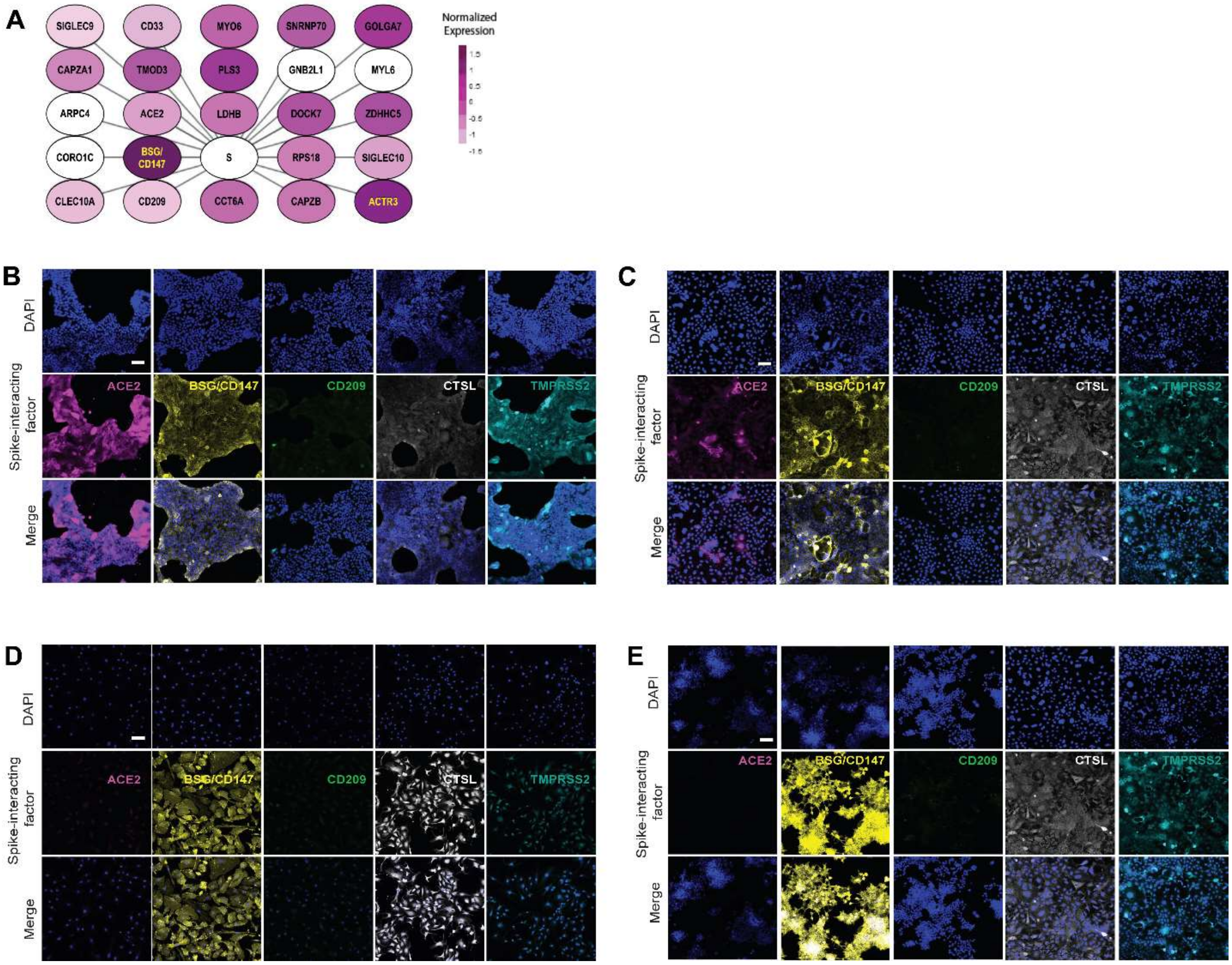


**Supplementary Figure 3**: (**A**) Chart showing twenty-four human proteins associated with spike-binding capabilities; S, Spike. BSG and ACTR3 are coloured yellow to improve legibility. SIGLEC9, Sialic acid-binding Ig-like lectin 9; CAPZA1, F-actin-capping protein subunit alpha-1; CLEC10A, C-type lectin domain family 10 member A; CD33, Myeloid cell surface antigen CD33; TMOD3, Tropomodulin-3; ACE2, Angiotensin

Converting Enzyme 2; BSG/CD147, Basigin/CD147 molecule; CD209, CD209 Antigen; MYO6, Unconventional myosin-VI; PLS3, Plastin-3; LDHB, L-lactate dehydrogenase B chain; GNB2L1/RACK, Receptor of activated protein C kinase 1; SNRNP70, U1 small nuclear ribonucleoprotein 70 kDa; DOCK7, Dedicator of cytokinesis protein 7**;** RPS18**,** 40S ribosomal protein S18; CAPZB, F-actin-capping protein subunit beta; GOLGA7, Golgin subfamily A member 7; ZDHHC5, Palmitoyltransferase SIGLEC10, Sialic acid-binding Ig-like lectin 10; ACTR3, Actin-related protein 3**;** MYL6, Myosin light polypeptide 6; CORO1C, Coronin-1C; ARPC4, Actin-related protein 2/3 complex subunit 4; CCT6A, T-complex protein 1 subunit zeta. (**B**-**E**) Immunocytochemistry analysis of ACE2, BSG/CD147, CD209, TMPRSS2 and CTSL expression, showing different levels of expression of the proteins in (**B**) Calu3 cells (**C**) Caco2 cells (**D**) Glomerular endothelial cells (gEndos) (**E**) HEK 293T cells.
